# What drives prey selection? Assessment of tiger food habits across the Terai-Arc landscape, India

**DOI:** 10.1101/2022.07.20.500750

**Authors:** Suvankar Biswas, Shrewshree Kumar, Meghna Bandhopadhyay, Shiv Kumari Patel, Salvador Lyngdoh, Bivash Pandav, Samrat Mondol

## Abstract

Large carnivores strongly shape ecological interactions within their respective ecosystems, but experience significant conflicts with human across their range due to their specific ecological resource requirements. The tiger (*Panthera tigris*) typifies the challenges faced by the large carnivore communities globally. India retains majority of the global tiger population with significant numbers outside protected areas (PAs) that are involved in conflicts from livestock predation and human/tiger death. To understand the prey selection patterns and spatio-temporal patterns of livestock predation-related conflict issues we investigated tiger food habits across the Indian part of the Terai-Arc Landscape (TAL), a globally-important tiger conservation landscape in India. We used 510 genetically-confirmed tiger faeces collected across the landscape and ascertained 10 wild and livestock as major prey species. Large-bodied species (sambar, swamp deer, nilgai, chital, wild pig and livestock) comprised ~94% of tiger diet, with sambar, chital and livestock were the major prey species. Habitat-specific (Shivalik-Bhabar and Terai) analyses show significantly different pattern of prey selections determined by abundance and prey body weight. Results also suggest that PA and non-PAs of Terai habitat are more prone to livestock depredation-related conflicts, and careful management interventions and community involvements are required to reduce such threats. We suggest long-term plans including population estimation of tigers and prey outside PAs, reducing grazing pressures and cattle enumeration, detailed investigation of tiger deaths etc. to ensure future tiger sustainability across this habitat.

## INTRODUCTION

Large carnivores play significant roles in maintaining ecological diversity and interactions within their respective biological communities (Steneck 2005). Being top predators, they impact the abundance of their prey species through predations (Ripple et al. 2014) and regulate the top-down cascade system (Hairston et al. 1960; del Rio et al. 2001; Treves and Karanth 2003). However, their ecological requirements of large, undisturbed habitats and adequate prey base (Linnell et al. 2001; Macdonald and Sillero-Zubiri 2002) often lead to competitions with humans (Inskip and Zimmermann 2009; Bhattarai and Fischer 2014) leading to loss of human, livestock and carnivore lives (Patterson et al. 2004; Inskip and Zimmermann 2009). Within large carnivores, the felines have particularly experienced severe reduction in their global population and geographic ranges (Ripple et al. 2014) mostly due to habitat loss, prey depletion, poaching and increase in negative human-carnivore interactions (Nowell 2000; Treves and Karanth 2003; Dinerstein et al. 2007; Chapron et al. 2008; Wikramanayake et al. 2010). Recent reports indicate that conflict arising from livestock predation by large felids is common across their distributions (for example, lion-Saberwal et al. 1994; Patterson et al. 2004; leopard-Zimmermann et al. 2005; Athreya et al. 2016; puma-Polisar et al. 2003; snow leopard-Bagchi and Mishra 2006; tiger-Chanchani et al. 2016; Chatterjee et al. 2017, etc.). As the predation patterns can be of species and area-specific nature, in-depth understanding of such patterns are important towards their conservation measures.

The tiger (*Panthera tigris*) exemplifies the problems faced by large feline predators globally as its range covers various habitat types such as evergreen forests, mangrove swamps, grasslands, tropical rainforests, low-land and hill forests, tropical and subtropical moist broadleaf forest and cold rocky-mountains (Sunquist 1999; Schaller 2009; Tilson and Nyhus 2009; Wikramanayake et al. 2010; Barber-Meyer et al. 2013). India currently retains ~80% of the world’s tigers (Jhala et al. 2008, 2010, 2015, 2020; Goodrich et al. 2015) and about one-third of these tigers live outside the protected areas (PAs) across different landscapes (Jhala et al. 2020) (possibly due to intra-species competition for resources) and involved in various levels of interactions with human (Gurung 2008; Dhanwatey et al. 2013). Such interactions often result in livestock depredation and at times loss of human lives, elevating negative attitude towards tiger conservation (Ogra and Badola 2008; Inskip and Zimmermann 2009; Goodrich 2010). Recent reports indicate a significant increase in human-tiger conflict in different parts of India (Karanth and Sunquist 1995; Bagchi et al. 2003; Andheria et al. 2007; Avinandan et al. 2008; Mondal et al. 2012; Chouksey and Singh 2018; Doubleday 2018; Sarkar et al. 2018; Ramesh et al. 2019; Bakhshi 2020), necessitating immediate attention and better understanding of this issue. In this regard, analyses of tiger food habits across large landscapes covering a mosaic of different habitat types could provide valuable insights towards spatio-temporal patterns of livestock predation-related conflict issues. Existing literature on tiger food habit are dominated by PA-centric data (Karanth and Sunquist 1995; Bagchi et al. 2003; Reddy et al. 2004; Andheria et al. 2007; Avinandan et al. 2008; Mondal et al. 2012; Sarkar et al. 2018), which are largely free from intense human activities (Biswas and Sankar 2002; Andheria et al. 2007; Upadhyaya et al. 2018), but scarce information is available from non-PAs and landscape scales. As landscape-scale studies of other long-ranging, large carnivores have already shown significant impacts of livestock predation in human-carnivore conflict (Ciucci et al. 1996; Hernández-SaintMartín et al. 2015; Lyngdoh et al. 2020), a study of tiger food habits in relation to their ecology, behaviour and other factors at large spatial scale would be of great conservation interest.

In this paper, tiger food habits were investigated across the Indian part of the Terai-Arc Landscape (TAL), a globally-important tiger conservation landscape in India. TAL consists of a mosaic of PAs and multiple use non-PAs covering multiple habitat types with varied distribution of large mammalian fauna (Johnsingh et al. 2004). This region has one of the highest human and livestock densities in India and has recently seen a ~34% increase in tiger population size (Jhala et al. 2020) and associated conflicts (Chanchani et al. 2014). The specific objectives were to (1) assess the tiger food habit in different habitat types across TAL (2) evaluate ecological variables that influence such patterns and (3) explore the patterns of livestock predation by tiger across TAL. These questions were addressed using field-collected tiger faecal samples across the entire landscape. In addition to understanding the food habit of tigers in different habitat types across a mosaic of PA and non-PAs in this landscape, we believe that these results have wider relevance for appropriate tiger-centric conservation plans within this rapidly-changing human-dominated landscape.

## MATERIALS AND METHODS

### Research permissions and ethical considerations

All required permissions for the surveys and biological sample collections were provided by the Forest Departments of Uttarakhand (Permit no: 90/5-6 978/6-32/56 and 3707/5-6), Uttar Pradesh (Permit no: 1891/23-2-12) and Bihar (Permit no: Wildlife-589). Due to non-invasive nature of sampling, no ethical clearance was required for this study.

### Study area

This study was conducted across ~15000 sq. km. forested habitat along the foothills of the Himalayas. This 900 km long linear stretch of habitat is known as the Terai-Arc Landscape (TAL) (Fig. 1) and recognised as one of the most productive habitats in the subcontinent (Macdonald and Loveridge 2010; Jhala et al. 2020), typically characterized by a mosaic of forests and grasslands covering both PAs and non-PAs. Overall, TAL represents three major habitat types namely Shivalik (hilly, undulating habitat of lower Himalayas in Uttarakhand and parts of northern Bihar), Bhabar (pebbly and bouldery foothills of lower Himalayas in Uttarakhand and parts of Bihar) and Terai (lowland region characterised by tall grasslands, scrub savannah, sal forests and swamps below the Himalayan foothills and north of Indo-Gangetic plains covering entire Uttar Pradesh, southern parts of Uttarakhand and Bihar) (Johnsingh et al. 2004) (Fig. 1, Supplementary data SD1). TAL also retains very high mammalian diversity including herbivores (gaur-*Bos gaurus*, nilgai-*Boselaphus tragocamelus*, sambar-*Rusa unicolor*, northern swamp deer-*Rucervus duvaucelii duvaucelii*, wild pig-*Sus scrofa*, chital-*Axis axis*, hog deer-*Hyelaphus porcinus*, barking deer-*Muntiacus muntjak*, goral-*Naemorhedus goral*, common langur-*Semnopithecus entellus*, rhesus macaque-*Macaca mulatta*, Tarai grey langur-*Semnopithecus hector* etc.), carnivores (tiger-*Panthera tigris*, leopard-*Panthera pardus*, wild dog-*Cuon alpines*, hyena-*Hyaena hyaena* etc.), and omnivores (sloth bear-*Melursus ursinus*, Asiatic black bear-*Ursus thibetanus* etc.) (Johnsingh et al. 2004; Jhala et al. 2020).

**Fig. 1:**
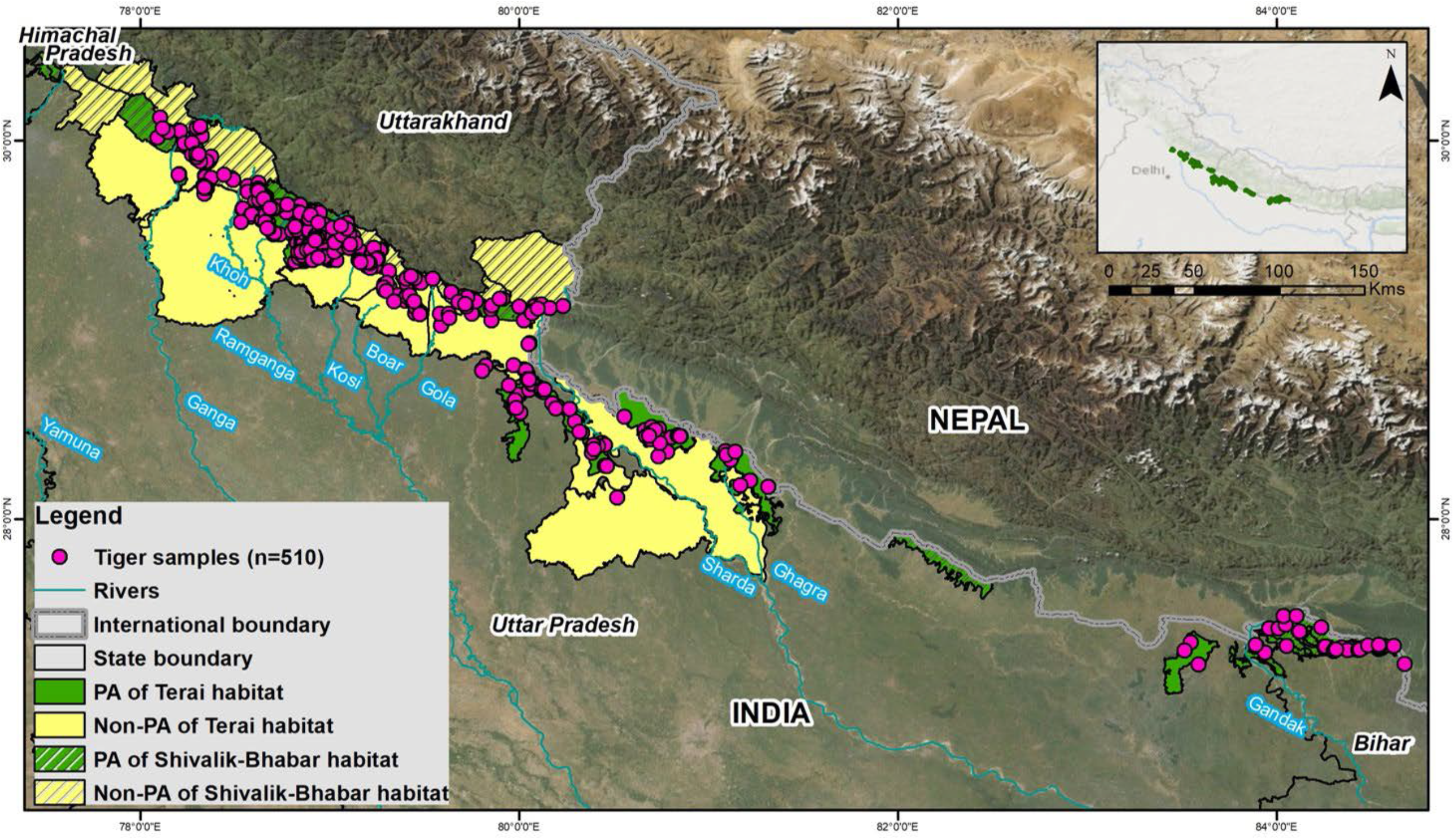
Distribution of tiger faecal samples (n=510) from Terai-Arc landscape (TAL), India used for food habit analyses in this study.

Latest assessment suggests that ~22% of India’s wild tiger population (n=646 (567-726)) is found across TAL (Jhala et al. 2020), living with one of the highest human and livestock densities in the subcontinent (Johnsingh et al. 2004). Interestingly, the densities of the tiger population and their potential prey species vary widely between the PAs/non-PAs, as the PAs generally receive better management interventions, protection and less human interferences when compared with the non-PAs (heavily disturbed by various anthropogenic activities) (Johnsingh et al. 2004; Chanchani et al. 2014). This study attempted to assess any possible difference in tiger food habit in TAL, as discerning such patterns have critical importance in habitat management in this region where tiger population are increasing along with human-tiger conflicts.

### Sample collection, species confirmation and prey identification

Field sampling was conducted across all the six tiger reserves (TRs) (Rajaji, Corbett, Amangarh, Pilibhit, Dudhwa and Valmiki), two wildlife sanctuaries (WLSs) (Nandhaur and Sohagibarwa), two conservation reserves (CRs) (Jhilmil Jheel and Pawalgarh) and 11 non-PAs (Haridwar, Lansdowne, Ramnagar, Terai West, Terai Central, Terai East, Haldwani, Champawat and South Kheri Forest Divisions (FDs) and Najibabad and Pilibhit Social Forestry Divisions (SFDs)) across the states of Uttarakhand, Uttar Pradesh and Bihar (Fig. 1, Supplementary data SD1). A total of 1689 large carnivore faecal samples were opportunistically collected (see Supplementary data SD2a) during extensive surveys between November 2014-December 2020 across TAL (Biswas et al. 2020). The research team surveyed the animal trails and other habitats and collected these samples in wax-papers, stored in dry boxes until shipped to the laboratory (Biswas et al. 2019), where they were finally stored in -20 °C freezer.

In the laboratory, the samples were first genetically ascertained through tiger-specific molecular markers (Mukherjee et al. 2007) to ensure only confirmed tiger samples were used to study the food habit. In brief, DNA was extracted from all field-collected large carnivore faecal samples using a modified Qiagen DNA extraction protocol (Biswas et al. 2019) and two tiger-specific mitochondrial DNA markers (Tig490F/R and Tig509F/R, Mukherjee et al. 2007) were used to ascertain tiger faeces. Tiger positive samples were selected for downstream food habit analyses and dried at 60°C for 72 hours in an oven (Unident Dental Electric Hot Air Oven, New Delhi, India). The undigested parts (hair, broken bones, hoof etc.) were separated by sieving the dried samples through sterile 0.5 mm stainless steel mesh. Permanent slides were prepared with the primary guard hairs to identify tiger prey species (Mukherjee et al. 1994) and were examined for cuticle, cortex and medulla structures (Mukherjee et al. 1994; Karanth and Sunquist 1995; Biswas and Sankar 2002; Avinandan et al. 2008; Bahuguna et al. 2010). To select the best approach for morphological identification of prey species, a pilot study with 45 tiger faeces was conducted where all three approaches (cuticular (for cuticle structure), cross-sectional (for cortex structure) and medullary methods (for medulla structure)) were used and the results were compared. Samples from Valmiki TR were used for this pilot study as this area retains the most varied prey assemblage in TAL (Johnsingh et al. 2004). Sample size estimation for diet analyses was conducted through a sample rarefaction curve (Hurlbert 1971; Heck Jr et al. 1975) in R (version 4.0.2) using the package vegan (Oksanen et al. 2013).

### Data analysis

#### Tiger food habit in different habitat types across TAL

During analyses, the sampled areas across TAL were stratified into different categories based on habitat types (Shivalik-Bhabar vs. Terai habitats) to comparatively ascertain any possible food habit differences. In each case, data on relative frequencies of occurrences (RFO) (Mukherjee et al. 1994), prey biomass (Andheria et al. 2007) and prey preference (Sankar et al. 2010) were calculated. For RFO calculation, formula i/j*100 was used, where ‘i’ represents the frequency of number of samples in which a specific prey occurs and ‘j’ represents the total frequency count of all prey species (Kruuk 1989; Mukherjee et al. 1994). To test any significant differences in RFOs of each prey species between and within Shivalik-Bhabar and Terai habitats, two-way ANOVA was performed as it can account for multiple groups (prey species) and factors (habitats). Additionally, ANOVA was used to test any significant differences in RFOs of major prey species between Shivalik-Bhabar and Terai. Prey biomass was calculated to overcome the issue of overestimation of small prey species in RFO analysis (Andheria et al. 2007; Chakrabarti et al. 2016) using Ackerman’s equation: Y=1.980 + 0.035X, where Y=weight of consumed prey in each faeces and X=mean body weight of a particular prey species (Ackerman et al. 1984; Karanth and Sunquist 1995). The mean body weight of prey was taken from Harihar et al. (2007); Harihar et al. (2011); Upadhyaya et al. (2018) (Table 1). The relative biomass of the prey (D) was calculated following Karanth and Sunquist (1995); Andheria et al. (2007) formula D= (A*Y)/∑ (A*Y)*100 where, A represents the RFO of each prey species. Two-way ANOVA was used to test any significant differences between and within the prey biomass of different habitats, whereas ANOVA was performed to test any significant differences in biomass of major prey species between Shivalik-Bhabar and Terai. Tiger prey preference in different habitats (Shivalik-Bhabar and Terai) was calculated using Ivlev’s electivity index: E=r-p/r+p, where, ‘r’ is the utilization of prey species or RFO and ‘p’ is the abundance or availability of the prey species in the area (Ivlev 1961). Its value ranges between -1 (no selection) to +1 (complete selection). Relative abundance index (RAI) for ungulates prey species from TAL has been used from (Jhala et al. 2020) (Table 2).

**Table 1:**
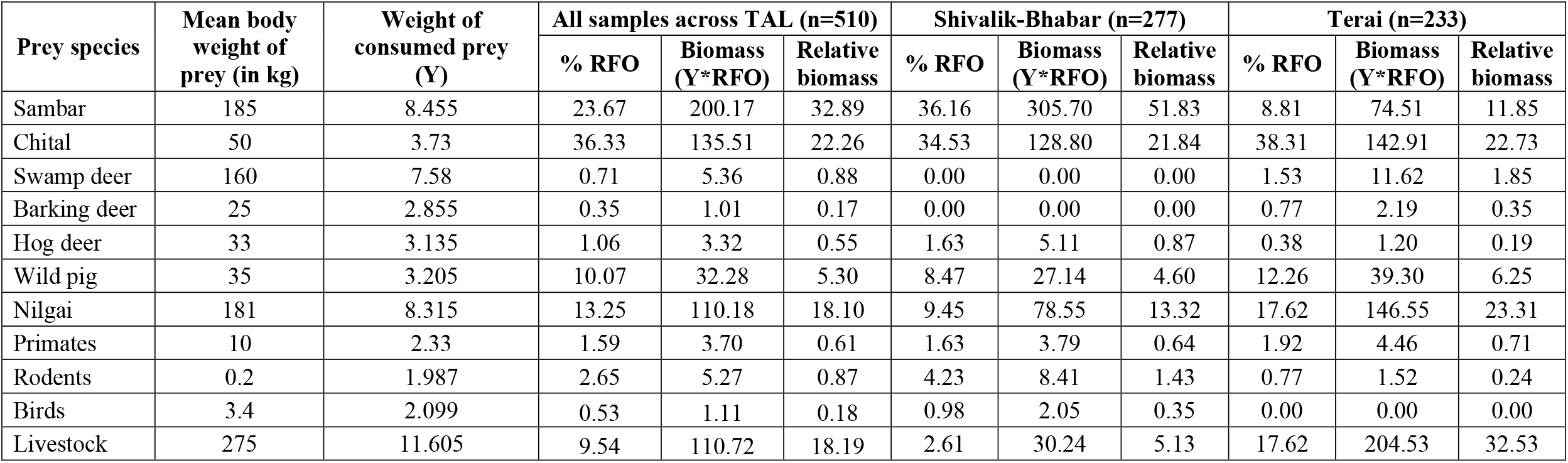
Details of the various tiger food habit parameters for all 11 prey species identified in this study. Results are presented for percentage of relative frequency of occurrence (% RFO), and biomass of the consumed prey species for entire TAL (n=510 samples), Shivalik-Bhabar (n=277 samples) and Terai habitats (n=233 samples), respectively.

| Prey species | Mean body weight of prey (in kg) | Weight of consumed prey (Y) | All samples across TAL (n=510) |  |  | Shivalik-Bhabar (n=277) |  |  | Terai (n=233) |  |  |
| --- | --- | --- | --- | --- | --- | --- | --- | --- | --- | --- | --- |
|  |  |  | % RFO | Biomass (Y*RFO) | Relative biomass | % RFO | Biomass (Y*RFO) | Relative biomass | % RFO | Biomass (Y*RFO) | Relative biomass |
| Sambar | 185 | 8.455 | 23.67 | 200.17 | 32.89 | 36.16 | 305.70 | 51.83 | 8.81 | 74.51 | 11.85 |
| Chital | 50 | 3.73 | 36.33 | 135.51 | 22.26 | 34.53 | 128.80 | 21.84 | 38.31 | 142.91 | 22.73 |
| Swamp deer | 160 | 7.58 | 0.71 | 5.36 | 0.88 | 0.00 | 0.00 | 0.00 | 1.53 | 11.62 | 1.85 |
| Barking deer | 25 | 2.855 | 0.35 | 1.01 | 0.17 | 0.00 | 0.00 | 0.00 | 0.77 | 2.19 | 0.35 |
| Hog deer | 33 | 3.135 | 1.06 | 3.32 | 0.55 | 1.63 | 5.11 | 0.87 | 0.38 | 1.20 | 0.19 |
| Wild pig | 35 | 3.205 | 10.07 | 32.28 | 5.30 | 8.47 | 27.14 | 4.60 | 12.26 | 39.30 | 6.25 |
| Nilgai | 181 | 8.315 | 13.25 | 110.18 | 18.10 | 9.45 | 78.55 | 13.32 | 17.62 | 146.55 | 23.31 |
| Primates | 10 | 2.33 | 1.59 | 3.70 | 0.61 | 1.63 | 3.79 | 0.64 | 1.92 | 4.46 | 0.71 |
| Rodents | 0.2 | 1.987 | 2.65 | 5.27 | 0.87 | 4.23 | 8.41 | 1.43 | 0.77 | 1.52 | 0.24 |
| Birds | 3.4 | 2.099 | 0.53 | 1.11 | 0.18 | 0.98 | 2.05 | 0.35 | 0.00 | 0.00 | 0.00 |
| Livestock | 275 | 11.605 | 9.54 | 110.72 | 18.19 | 2.61 | 30.24 | 5.13 | 17.62 | 204.53 | 32.53 |

**Table 2:** Details of Tiger prey preferences (Ivlev’s index) within protected areas (PAs) at different scales considering all prey species (n=11). Results are presented for the PAs of entire TAL, Shivalik-Bhabar and Terai habitats, respectively.

| PA-TAL (n=308) |  |  |  | PA-Shivalik-Bhabar (n=184) |  |  | PA-Terai (n=124) |  |  |
| --- | --- | --- | --- | --- | --- | --- | --- | --- | --- |
| Species | Utilisation of prey species (r) | Abundance of prey species<sup>a</sup> (p) | Ivlev's index | Utilisation of prey species (r) | Abundance of prey species<sup>a</sup> (p) | Ivlev's index | Utilisation of prey species (r) | Abundance of prey species<sup>a</sup> (p) | Ivlev's index |
| Sambar | 21.10 | 18.41 | 0.40 | 32.34 | 30.27 | 0.52 | 5.52 | 0.92 | -0.27 |
| Chital | 39.02 | 36.82 | 0.62 | 39.80 | 27.72 | 0.59 | 37.93 | 33.84 | 0.59 |
| Swamp deer | 1.16 | 0.36 | -0.77 | 0.00 | 0.00 | -1.00 | 2.76 | 1.26 | -0.56 |
| Barking deer | 0.58 | 0.72 | -0.88 | 0.00 | 10.19 | -1.00 | 1.38 | 1.76 | -0.75 |
| Hog deer | 1.45 | 0.72 | -0.73 | 2.49 | 0.16 | -0.61 | 0.00 | 5.58 | -1.00 |
| Wild pig | 11.85 | 13.00 | 0.13 | 9.45 | 9.16 | -0.04 | 15.17 | 10.70 | 0.22 |
| Nilgai | 13.01 | 16.25 | 0.18 | 8.46 | 2.15 | -0.10 | 19.31 | 6.29 | 0.33 |
| Primates | 1.73 | 2.17 | -0.68 | 1.49 | 4.87 | -0.75 | 2.07 | 10.55 | -0.65 |
| Rodents | 1.73 | 1.81 | -0.68 | 1.99 | 1.05 | -0.68 | 1.38 | 1.13 | -0.75 |
| Birds | 0.58 | 0.36 | -0.88 | 1.00 | 9.98 | -0.82 | 0.00 | 12.46 | -1.00 |
| Livestock | 7.51 | 9.39 | -0.09 | 2.49 | 17.46 | -0.61 | 14.48 | 21.70 | 0.20 |

<sup>a</sup>data taken from Jhala et al. 2020

#### The relationship between tiger food habit with ecological variables

To ascertain any relationship of tiger food habit (RFOs and prey biomass (response variables)) with ecological variables (prey abundance and prey body weight (explanatory variables) of prey species) in Shivalik-Bhabar and Terai, general linear model (GLM) was performed in R using package lme4 (Bates et al. 2015). The prey RFOs and biomass were run separately against the explanatory variables for both habitats, respectively. Model parameterization was performed to estimate the effect size of both the habitats (Kéry and Royle 2020) as a response to the explanatory variables. Collinearity between explanatory variables was checked as it might reduce the precision of the estimated coefficients. R-square value (0=low to 1=high) was used to detect collinearity. The test was performed in R platform using package lattice (Sarkar 2018).

Finally, to visualize the overall prey species predation pattern by tiger in Shivalik-Bhabar and Terai (species grouping in different habitat), Non-metric Multidimensional Scaling (NMDS) (Clarke 1993) ordination was used. RFO, biomass and abundance of prey species in Shivalik-Bhabar and Terai habitats and their respective body weights were used as variables for the analysis. NMDS was performed in R through meta MDS function using vegan package (Oksanen et al. 2013). The models (containing all the variables) were run over different dimensions (1-3) with Bray-Curtis distance and the best model was selected based on the lowest stress value (Zuur et al. 2007).

#### Patterns of livestock predation by tiger in TAL

The TAL region has experienced a 34% increase in tiger numbers in last four years (Jhala et al. 2020) and increase in human-tiger conflict driven by livestock predation and resulting retaliatory killing of tigers (Chanchani et al. 2014). Kernel density estimation (KDE) technique was used to assess the livestock predation hotspots across TAL, as this approach smoothens discrete point data into a continuous surface area (Silverman 1986; Hart and Zandbergen 2014). As recent reports suggest much-restricted pattern of such conflict incidences in the Terai habitat (possibly due to much more human and livestock densities than in the Shivalik-Bhabar habitat) (Chanchani et al. 2014), only samples from this region were further categorised (PA-three TRs and one WLS and non-PA-five FDs and two SFDs; Fig. 1, Supplementary data SD1) to assess the patterns of livestock prey presence. Analysis of similarities (ANOSIM) was performed using Bray-Curtis distance in R using package vegan (Oksanen et al. 2013). The R statistic (ratio of dissimilarities in observations within a group to that of dissimilarities between groups) was used to understand the similarity of livestock predation rate by tigers. The closer the R statistic value to 0, more the similarity of observations between two groups.

## RESULTS

A total of 525 genetically confirmed tiger faecal samples from TAL were used in this study (Supplementary data SD2b). Out of these tiger faeces, 510 samples (success rate - 97.1%) produced data on the prey species assemblage. The remaining samples (n=15, 2.9%) contained damaged hairs for accurate species identification and were discarded from analyses. The comparison among three morphological prey identification methods (cuticular, cross sectional and medullary) (Supplementary data SD3, Supplementary data SD4) on 45 tiger faeces from Valmiki TR showed variable success rates in species identification. The medullary method showed 100% success rate with this set of faeces, whereas the other two methods (cuticular and cross-sectional) achieved only ~40% success rates, respectively. Based on the results, the medullary method was subsequently used with the remaining samples for tiger prey identification.

Tiger consumed mainly five large (sambar, swamp deer, nilgai, wild pig and chital) and five relatively small bodied (hog deer, barking deer, primates, rodents and birds) wild ungulate species along with livestock (large-bodied domestic ungulate). These six large-bodied (wild and domestic) prey species comprised 93.7% (RFO) of tiger diet whereas small-bodied prey species contributed only 6.3% (RFO). RFO of chital (36.3%) was found to be the highest followed by sambar (23.6%) (Table 1). However, biomass of sambar (32.9%) was highest followed by chital (22.3%) (Table 1). Among all the 11 identified prey species, chital (Ivlev’s value=0.62) was the most preferred followed by sambar (Ivlev’s value=0.4) (Fig. 2A, Table 2). The rarefaction curve flattened beyond 280 samples and no new prey species was identified (Supplementary data SD5).

**Fig. 2:**
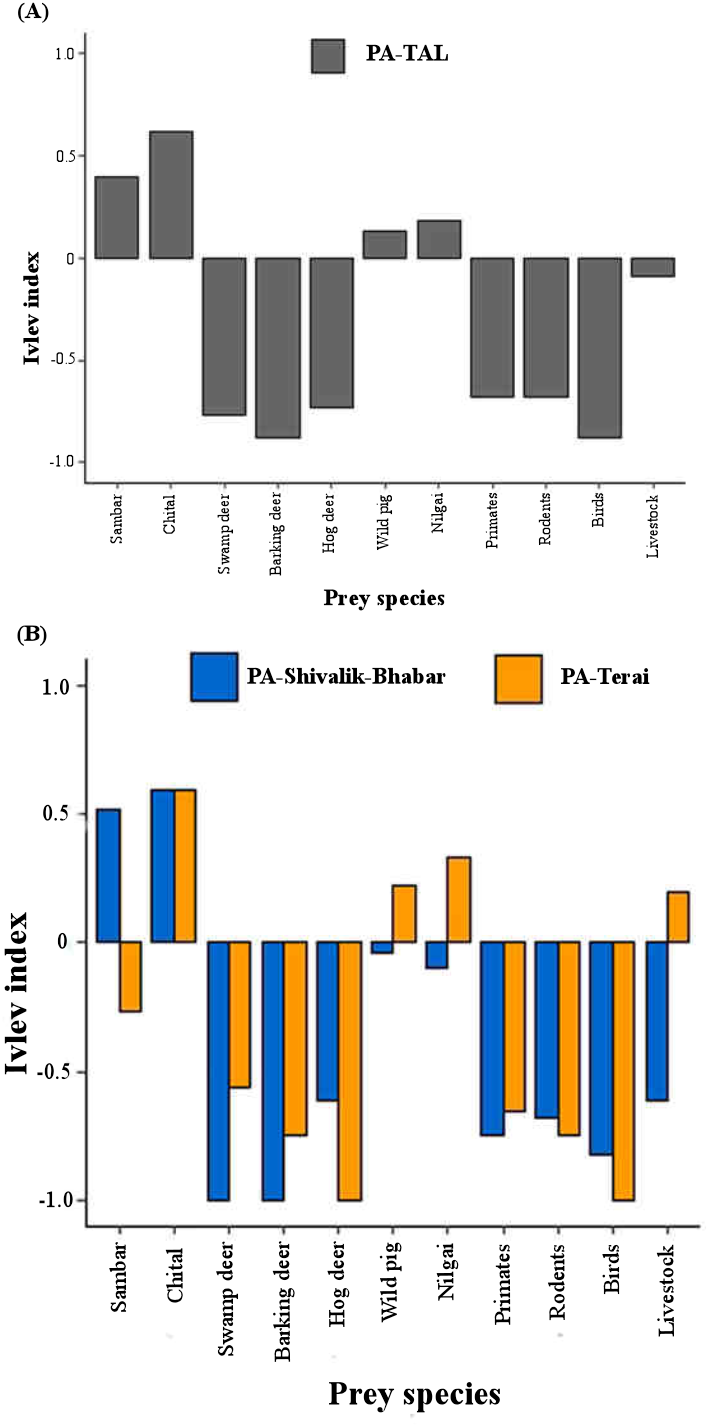
Tiger prey preferences (Ivlev’s index) within protected areas (PAs) at different scales. The top panel (A) indicate preferences at the entire TAL and the bottom (B) panel shows preference patterns at Shivalik-Bhabar and Terai habitats separately. The major differences in prey preference between these two habitats were for large-bodied species sambar, livestock, wild pig and nilgai.

### Tiger food habit in different habitat types across TAL

A total of 307 and 261 prey occurrences were found from Shivalik-Bhabar (n=277 samples) and Terai (n=233 samples) habitats, respectively. Majority of the samples (~89%) (Shivalik-Bhabar-247 samples (89.2%) and Terai-206 samples (88.4%)) contained single prey species, whereas only ~11% of the samples (n=30 (10.8%) in Shivalik-Bhabar and n=27 (11.6%) in Terai) had multiple prey species. In Shivalik-Bhabar, a total of nine prey species were identified: four large-bodied wild prey (sambar, chital, nilgai and wild pig), four relatively small-bodied wild prey (rodents, primates, birds and hog deer) and livestock. However, in Terai along with all the nine prey species mentioned above, two additional species swamp deer (large-bodied prey) and barking deer (small-bodied prey) were also found. In Shivalik-Bhabar, sambar RFO (36.2%) and biomass (51.8%) were found to be the highest whereas chital RFO (38.31%) and livestock biomass (32.5%) were highest in Terai habitat. Chital was most preferred prey in both the habitats (Shivalik-Bhabar=0.59, Terai=0.59) (Fig. 2B, Table 2), followed by sambar (Shivalik-Bhabar=0.52) and nilgai (Terai=0.33) (Fig. 2B, Table 2). The two-way ANOVA analyses with all 11 prey species between two habitat types showed no significant differences in both RFO and biomass. However, overall comparison within habitats indicated significant differences in RFO (p-value=0.01, alpha (α)=0.05, F statistic=4.89) but no differences in biomass. Further comparative analyses of only major prey species (n=6) between habitats (ANOVA) showed contrasting results, where RFO and biomass of sambar (p-value<0.001, alpha (α)=0.05, F statistic=58.3) and livestock (p-value=0.01, alpha (α)=0.05, F statistic=7.3) showed significant differences, but chital, wild pig and nilgai were not different.

### The relationship between tiger food habit with ecological variables

The GLM analyses indicate that both RFO (AIC=152.8, p-value<0.01, R2 adjusted=0.74) and biomass AIC=156.1, p-value<0.01, R2 adjusted=0.73) have significant positive relationship to prey abundance in Shivalik-Bhabar (t-value=3.6, p<0.01) and Terai (t-value=3.8, p<0.01) habitats. However, these two habitats showed contrasting relationship patterns with prey body weight, where no significant relationship was seen between RFO and body weight in either of the habitats but it was positively related with prey biomass in Terai (t-value=3.1, p<0.01) (Fig. 3, Table 3). The R-square values of collinearity test between prey abundance and body weight were 0.41 and 0 for Shivalik-Bhabar and Terai, respectively, indicating that the explanatory variables were not correlated in either of the habitats.

**Table 3:** Results showing the relationships between the ecological variables (habitat type and prey abundance) and food habit (RFO and prey biomass) based on GLM analyses.

| S.No. | Response variables | Explanatory variables | Estimate | Standard error | t value | Pr(>\|t\|) |
| --- | --- | --- | --- | --- | --- | --- |
| 1 | Prey RFO | Shivalik-Bhabar habitat | -1.08 | 3.87 | -0.28 | 0.78 |
| 2 | Prey RFO | Terai habitat | -2.69 | 3.93 | -0.69 | 0.50 |
| 3 | Prey RFO | Shivalik-Bhabar habitat: Prey abundance | 0.98 | 0.27 | 3.65 | 0.00 |
| 4 | Prey RFO | Terai habitat: Prey abundance | 0.92 | 0.24 | 3.81 | 0.00 |
| 5 | Prey RFO | Shivalik-Bhabar habitat: Prey body weight | 0.00 | 0.03 | 0.02 | 0.99 |
| 6 | Prey RFO | Terai habitat: Prey body weight | 0.03 | 0.03 | 1.27 | 0.22 |
| 7 | Prey biomass | Shivalik-Bhabar habitat | -3.92 | 4.18 | -0.94 | 0.36 |
| 8 | Prey biomass | Terai habitat | -4.80 | 4.25 | -1.13 | 0.28 |
| 9 | Prey biomass | Shivalik-Bhabar habitat: Prey abundance | 0.92 | 0.29 | 3.19 | 0.01 |
| 10 | Prey biomass | Terai habitat: Prey abundance | 0.66 | 0.26 | 2.50 | 0.02 |
| 11 | Prey biomass | Shivalik-Bhabar habitat: Prey body weight | 0.04 | 0.03 | 1.39 | 0.19 |
| 12 | Prey biomass | Terai habitat: Prey body weight | 0.09 | 0.03 | 3.10 | 0.01 |

**Fig. 3:**
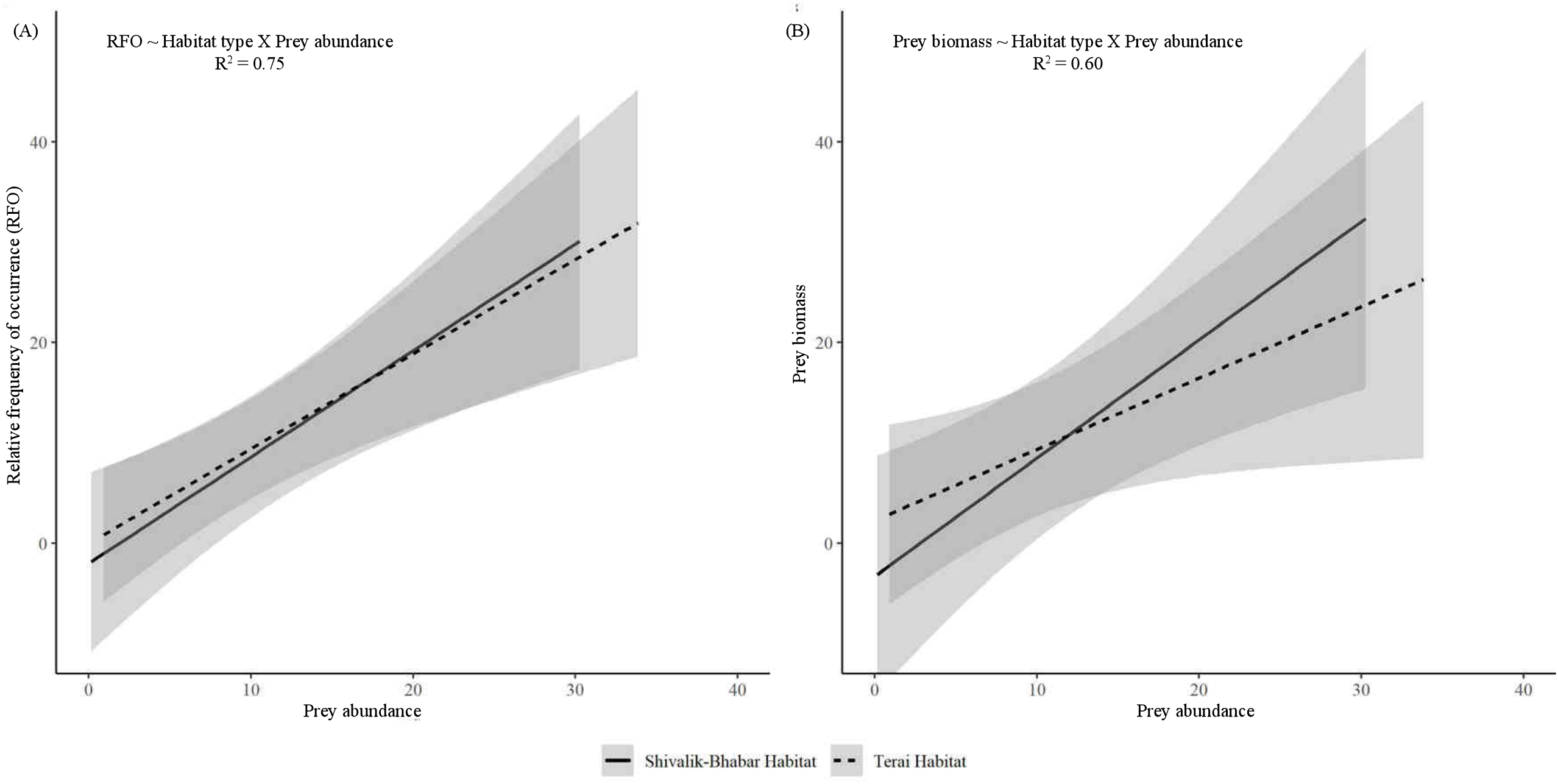
Results of the GLM analyses showing the relationships between ecological variables (habitat type and prey abundance) and dietary pattern (RFO and prey biomass). Both left pane (A, with RFO) and right pane (B, with prey biomass) shows positive relation to abundance and habitat type.

The two-dimensional NMDS ordination results indicated that sambar is the most dominant prey species (based on RFO, biomass and abundance) in Shivalik-Bhabar habitat whereas livestock was found to be the dominant prey species (supported by RFO, biomass, abundance and body weight) in Terai habitat (stress value=0.024, R2 value=0.99) (Fig. 4). Nilgai, wild pig and chital showed similar predation rates, biomass and abundance in both the habitats whereas swamp deer, hog deer, primates, birds and barking deer did not show any specific predation pattern. The Shepherd diagrams of all the models are provided (Supplementary data SD6).

**Fig. 4:**
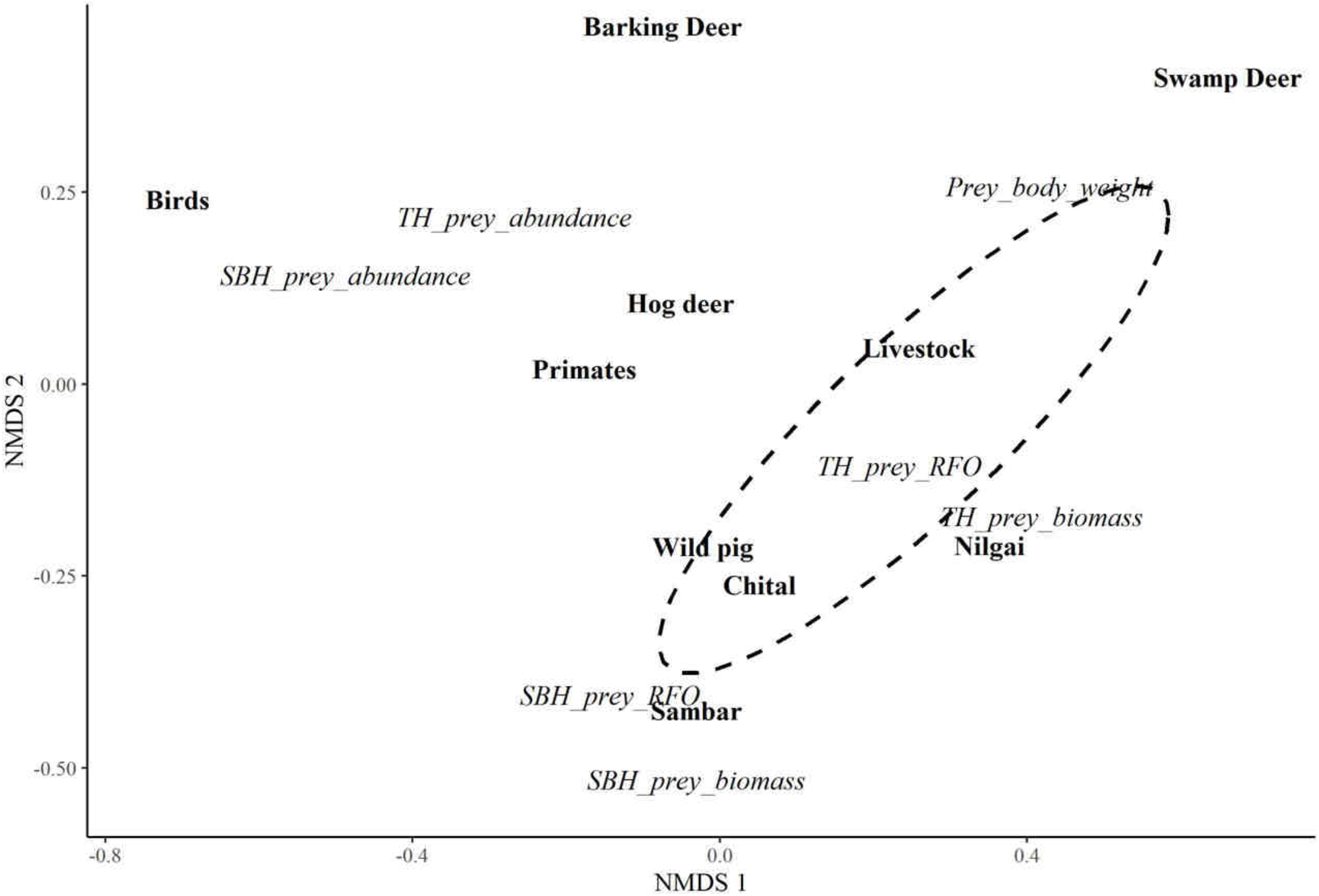
Two-dimensional NMDS plot depicting large-bodied species are the most dominant prey in TAL. While sambar is the dominant prey in Shivalik-Bhabar habitat (SBH), livestock, chital, nilgai and wild pig are the major prey species in Terai habitat (TH).

### Patterns of livestock predation by tiger in TAL

Out of all the processed samples (n=510), ~10% tiger faeces (n=53) contained livestock evidence, mostly found in the Terai habitat (n=46, ~87%). The KDE hotspot mapping corroborated this pattern (Fig. 5) where Amangarh TR, Terai West FD and south-eastern boundary of Corbett TR were found as major livestock predation hotspots and all of them are in Terai habitat (Fig. 5). Further, Najibabad SFD, Terai Central and East FDs, Pilibhit and Dudhwa TRs of Terai habitat were also identified as hotspots (Fig. 5). Close stratification of the livestock-containing faeces showed that ~46% (n=21) of these samples were collected from the PAs in Terai. This pattern was also confirmed by ANOSIM analysis of RFO between the Terai PA vs. non-PAs indicating similar predation rates of livestock between them (R statistic=0.40, p-value=0.026).

**Fig. 5:**
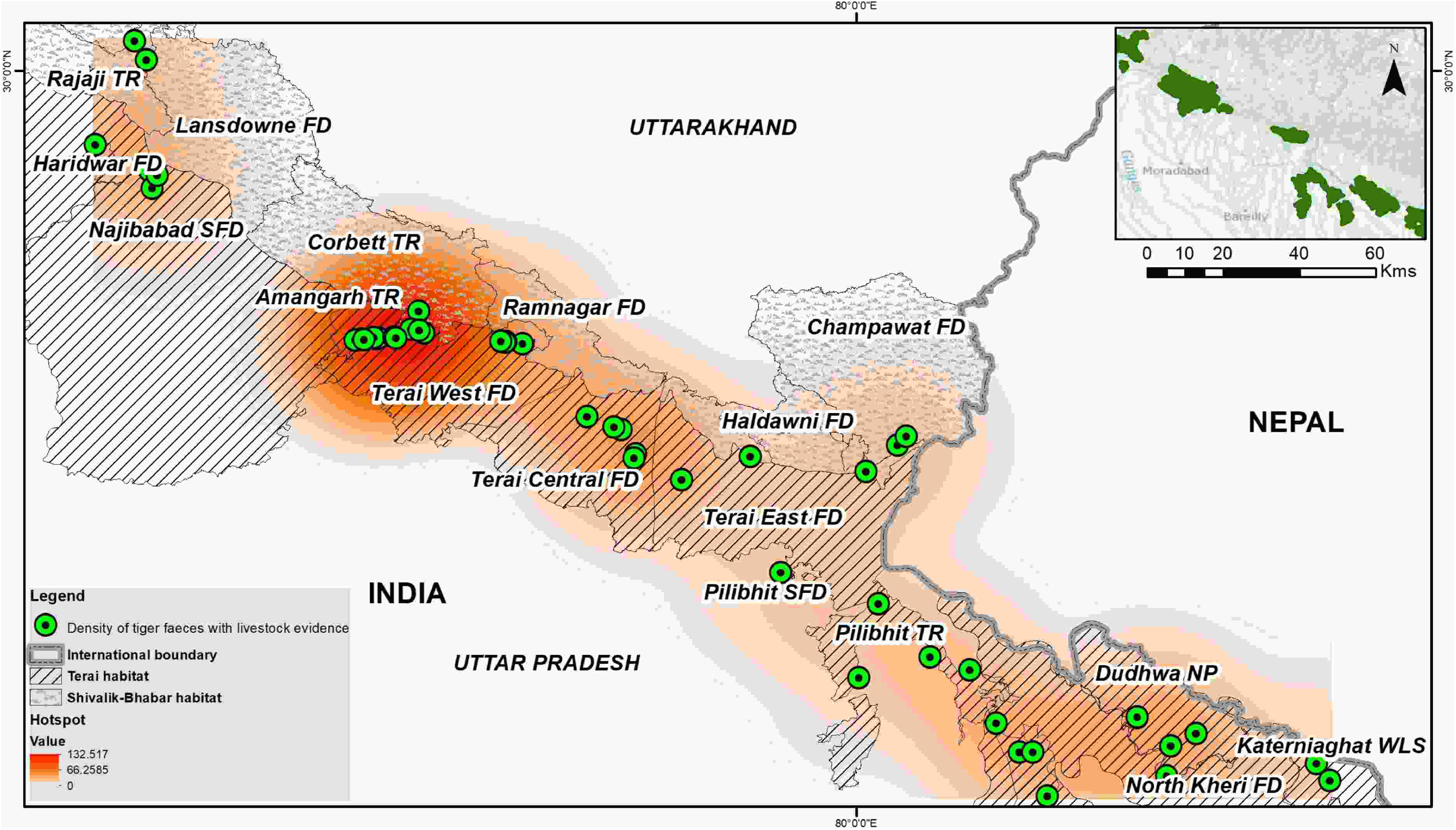
Identification of major livestock depredation hotspot areas in the Terai habitat through Kernel density estimation (KDE) approach. The results indicate a major hotspot around the south-east part of Corbett Tiger Reserve (TR), Amangarh TR and Terai West Forest Division (FD) followed by Najibabad Social Forestry Division (SFD), Terai Central and East FDs, Pilibhit and Dudhwa TRs.

## DISCUSSION

Most of the existing studies have focused on food habits at PAs (Bagchi et al. 2003; Avinandan et al. 2008; Kapfer et al. 2011; Kolipaka et al. 2017; Upadhyaya et al. 2018) due to their forest-centric habitat preferences (Smith et al. 1998; Karanth 2003). Limited insights on food habits at landscape levels (including non-PAs) are available. We found habitat-level ecological variables (prey abundance, prey body weight) influence tiger dietary patterns in a mosaic of PAs and non-PAs habitats. The methods used in this study are comparable to earlier research in India as well as other places for tiger dietary analyses (molecular species identification, hair medullary pattern etc.) (Andheria et al. 2007; Avinandan et al. 2008; Upadhyaya et al. 2018). The comparative analyses of the different morphological approaches (cuticular, medullary and cross-section) for prey identification confirmed the efficacy of the medullary approach over the others (Dharaiya and Soni 2012; Souza and Azevedo 2020) (Supplementary data SD3, Supplementary data SD4). Given that the Indian subcontinent currently retains the largest tiger population across the range residing within the most varied habitat types (Jhala et al. 2020, 2021), such information will be of great importance towards habitat management and conflict mitigation. However, it is important to point out that this study did not attempt to cover any potential seasonal differences in dietary patterns, and future efforts should focus on this aspect.

As expected, the results of this study showed that large-bodied prey constitute most of the tiger diet (94%), corroborating with majority of the existing studies (Sunquist 1981; Biswas and Sankar 2002; Hayward et al. 2014; Chatterjee et al. 2017). Within the Indian part of TAL, earlier studies have reported sambar, chital, wild pig and nilgai as the major prey species (Harihar et al. 2007, 2009; Basak et al. 2016) along with four-horned antelope and gaur from Nepal part of TAL (Bhandari et al. 2017; Thapa and Kelly 2017; Upadhyaya et al. 2018). Taken together, it can be inferred that the above-mentioned ungulate species are the major prey for tiger across TAL (both India and Nepal). Livestock also significantly contributed in tiger diet (Fig. 2A), possibly due to their overlapping distribution with tigers across this human-dominated landscape (Patterson et al. 2004; Harihar and Pandav 2012; Basak et al. 2016; Chanchani et al. 2016; Chatterjee et al. 2017). Overall, the data suggests that sambar, chital and livestock are the most predated prey species across TAL. However, when the data was stratified certain habitat-specific patterns were observed. For example, in Shivalik-Bhabar habitat sambar was the most predated (inferred from RFO values) and consumed (inferred from biomass) prey species, whereas in Terai chital was the most predated (based on RFO) and livestock was most consumed (based on biomass) prey, respectively. While earlier studies support this pattern from respective areas (Shivalik-Bhabar-Harihar et al. 2011, Terai-Basak et al. 2016), this landscape level pattern elucidated the underlying ecological factors that determines tiger prey selection across different habitats. The model-based correlations further substantiated these relationships where the results indicated that tiger prey predation and consumption is governed by abundance in Shivalik-Bhabar (RFO vs prey abundance 0.98 ± 0.27, p<0.01; prey biomass vs prey abundance 0.92 ± 0.29, p=0.01). Sambar is known to be present in higher density in the undulating terrains of Shivalik-Bhabar habitat (Johnsingh et al. 2004; Jhala et al. 2015) and therefore became the major prey species. On the other hand, the Terai habitat showed a positive relationship between predation and abundance (similar to Shivalik-Bhabar) but prey consumption was related to both abundance and body weight (RFO vs prey abundance 0.92 ± 0.24, p<0.01; prey biomass vs prey body weight 0.09 ± 0.03, p=0.01). This can be explained by the fact that Terai habitat lacks large-bodied wild ungulate species (such as sambar) (Johnsingh et al. 2004) and therefore tigers prefer highly abundant chital as prey. However, livestock being large-bodied prey (high biomass) was also consumed by tiger. Such difference in prey selection pattern was further corroborated by the multivariate NMDS analyses showing sambar and livestock as dominant prey species in Shivalik-Bhabar and Terai habitats, respectively. Combined together, we can infer that tiger prey selection was governed by prey body weight (as suggested by (Mondal et al. 2012; Upadhyaya et al. 2018; Krishnakumar et al. 2022) as large-bodied prey (sambar and livestock) was consumed despite their relatively low abundances in their respective habitats.

Our analyses of the livestock predation data showed much higher intensity of such incidences in Terai habitat (RFO= 17.62%) than Shivalik-Bhabar (RFO= 2.61%), possibly due to higher livestock abundance in Terai (Harihar and Pandav 2012; Malviya and Ramesh 2015; Chanchani et al. 2016; Anwar and Borah 2020; Bakhshi 2020). The results identified major livestock predation hotspot in the tri-junction of Corbett TR, Amangarh TR and Terai West FD, an area that has earlier been reported to have high numbers of livestock depredation events (Bargali and Ahmed 2018). Apart from the diet-based information presented here earlier studies also reported high livestock predation in the fringe areas of Pilibhit TR and Dudhwa NP (Terai habitat) (Basak et al. 2016; Chatterjee et al. 2017). Such patterns are probably expected as most of these areas are part of tiger source-recipient populations (Corbett, Pilibhit and Dudhwa TR are known sources and Amangarh TR and Terai West FD are recipient populations) (Biswas et al. 2020), where dispersing tigers get easy access to livestock/feral cattle (Chatterjee et al. 2017; Bargali and Ahmed 2018). One of the interesting pattern observed in this study was similar livestock predation rates (r value=0.4 from ANOSIM analysis) in PAs and non-PAs (7.5% and 12.6% RFOs, respectively), strongly indicating presence of livestock within the PAs. This would be a major conservation concern in this landscape as such events often lead to human-tiger conflict (Bagchi et al. 2003; Harihar et al. 2009; Chatterjee et al. 2017; Bargali and Ahmed 2018; Sarkar et al. 2018). Chatterjee et al. (2017) has reported that poor husbandry practices has led to higher cattle density within the forest region in some parts of Terai habitat. Careful management interventions in the form of better husbandry practices (stall feeding), reducing grazing pressure from inside PAs, adequate compensation programs in a timely manner etc. can reduce such conflict threats (Azevedo and Murray 2007; Malviya and Ramesh 2015; Bakhshi 2020; Badola et al. 2021). Successful implementation of such practices have reduced human-animal conflicts in different parts of the globe (for example, Jaguar-Quigley and Crawshaw Jr, 1992; Hoogesteijn et al. 2002; Azevedo and Murray 2007; Navarro-Serment et al. 2011; Cavalcanti et al. 2012; Hyena-Yirga and Bauer 2010; Sintayehu et al. 2016; Lion-Meena et al. 2014; Cheetah-Selebatso et al. 2008; Marker and Boast 2015; Leopard-Athreya and Belsare 2007). Although the Terai habitat retains high wild prey densities within the Pas (Johnsingh et al. 2004; Harihar et al. 2009; Chanchani et al. 2014; Jhala et al. 2020) reduction of human-tiger conflicts in the non-PAs (being used by transient tigers) would be critical for long-term tiger population sustainability across this habitat.

## CONSERVATION IMPLICATIONS

The presence of tiger faeces across the landscape, tiger food habits and patterns of habitat-specific prey selection direct us towards certain critical management interventions in this landscape. For example, presence of large numbers of tiger faeces in non-PAs confirms much extensive distribution of the species in TAL. However, current tiger monitoring and population estimation activities conducted by National Tiger Conservation Authority, Government of India are mostly limited to only PAs and selected non-PAs (Jhala et al. 2020), and we feel that expanding such efforts to outside PAs (particularly in the identified corridors, FDs, SFDs and CRs) would be very helpful in understanding their population patterns in these areas and help in developing better management plans. This will also help in getting substantial information on wild prey species in these areas. Further, the results also indicate that certain PAs and other area in Terai habitat are potential conflict hotspots due to livestock depredation events (Malviya and Ramesh 2015; Bargali and Ahmed 2018). Significant proportion of these evidences are from inside PAs indicating presence of livestock within them. Proper monitoring of tiger and prey populations (through active patrolling) and community-driven plans are required to assess the density of livestock/feral cattle within tiger habitats so necessary management interventions can be prepared. Once implemented, they can reduce tiger-human conflict and probability of disease transmission from domestic ungulates to wildlife (Martin et al. 2011; Khanyari et al. 2021). Finally, detailed investigation of any tiger death across India (both natural and retaliatory killings) needs to be conducted and records need to be examined thoroughly (as they had shown to reduce up to 50% decrease in population size in Russian far east, Miquelle et al. 2006), to ascertain causal reasons for appropriate management implications.

The TAL region is considered one of the globally important tiger conservation landscape that has shown significant increasing trend in population size (~34%) in the last decade (Jhala et al. 2015, 2020). Given that this expanding population competes for habitat with very high human density, the conservation plans require intensive effort to manage the remaining habitats and reduce human-tiger interactions.

## ACKNOWLEDGEMENTS

We acknowledge the Forest Departments of Uttarakhand, Uttar Pradesh and Bihar for research permits and logistic support during field work. Our sincere thanks to the Director, Dean, Research Coordinator, Laboratory in-charge and Nodal Officer, WFCG Cell for their continuous support. This work would not have been possible without the support from Harshvardhan S. Rathod, Annu, Bura, Abbhi, Ranjhu, Inam and Imam during fieldwork. We greatly appreciate guidance from Dr. S. P. Goyal and Mr. Vinod Thakur and technical supports from A. Madhanraj, Supriya Bhatt (DNA work) and Rakesh Sundriyal and Chitrapal (food habit analyses).

## AUTHOR CONTRIBUTIONS

SM, BP, SL and SB conceived the study design. SB, SKP and BP conducted field data collection. Molecular work was performed by SB and SKP and prey identification data was generated by SK. Data analysis was done by SK and MB. SB, SK, MB and SM wrote the first draft and all authors contributed in the final version for submission.

## SUPPLEMENTARY DATA

**Supplementary data SD1:** Details of the sampling sites, their characteristics and sample size (area-wise) in TAL.

**Supplementary data SD2:** Map of the study area showing the (A) distribution of total number of field-collected faecal samples (n=1689); and (B) distribution of genetically confirmed tiger faecal samples (n=525).

**Supplementary data SD3:** Pictorial representation of different prey hair identification approaches (cuticular, medullary and cross-sectional) used in this study. Results are presented for all 11 prey species identified in tiger food habit.

**Supplementary data SD4:** Comparison among different prey hair identification approaches (cuticular, medullary and cross-sectional) used in this study.

**Supplementary data SD5:** The rarefaction curve indicating sample sufficiency for tiger food habit assessment.

**Supplementary data SD6:** The Shephard diagrams of all the models tested in NMDS analyses.

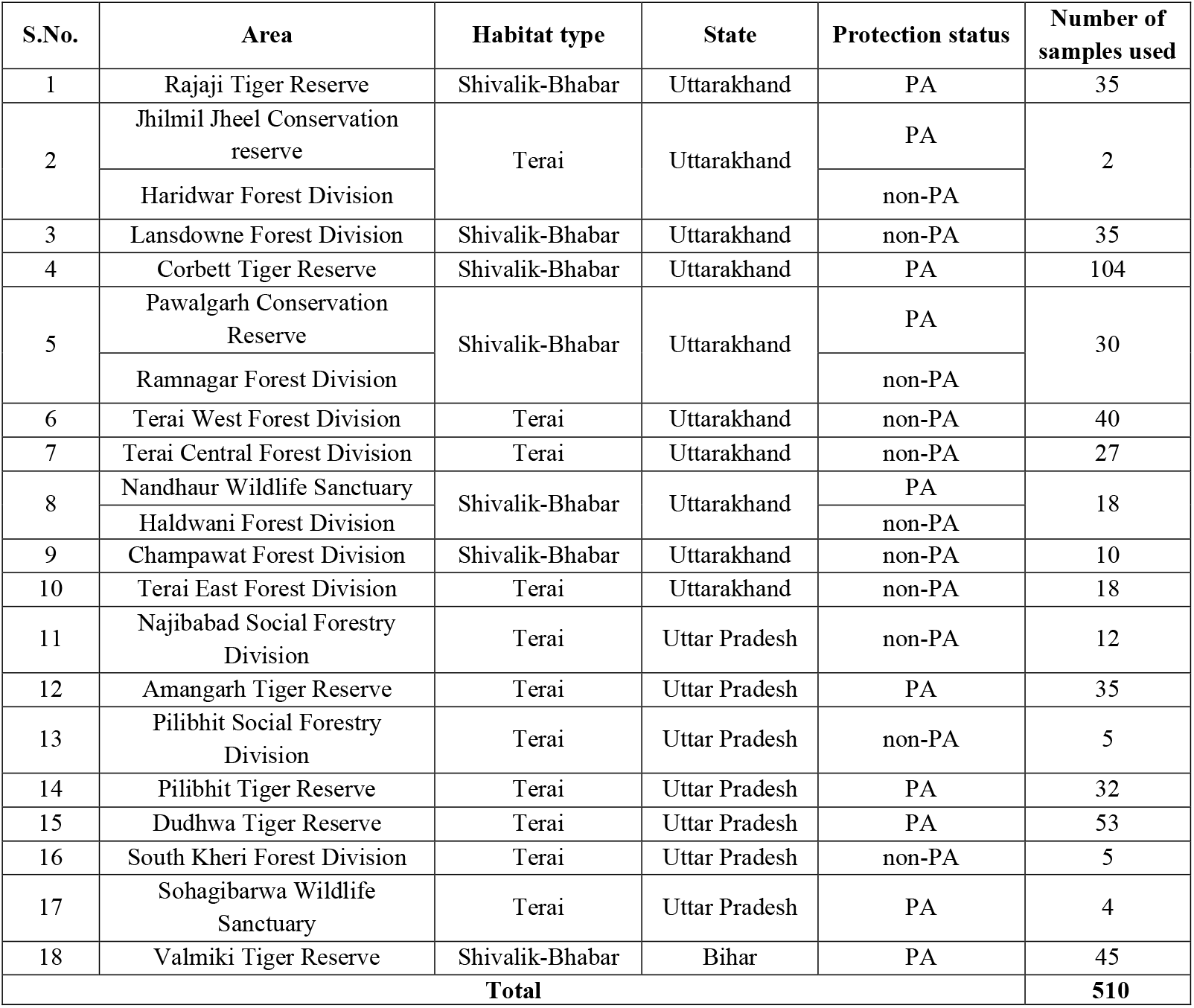

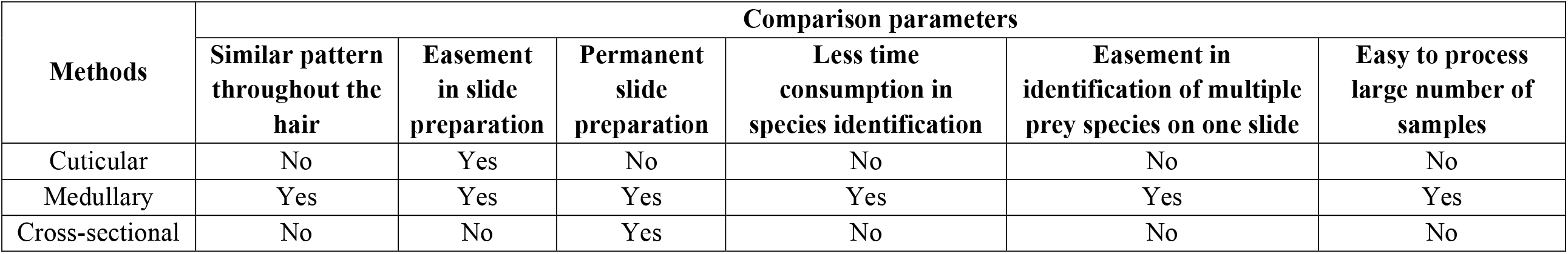

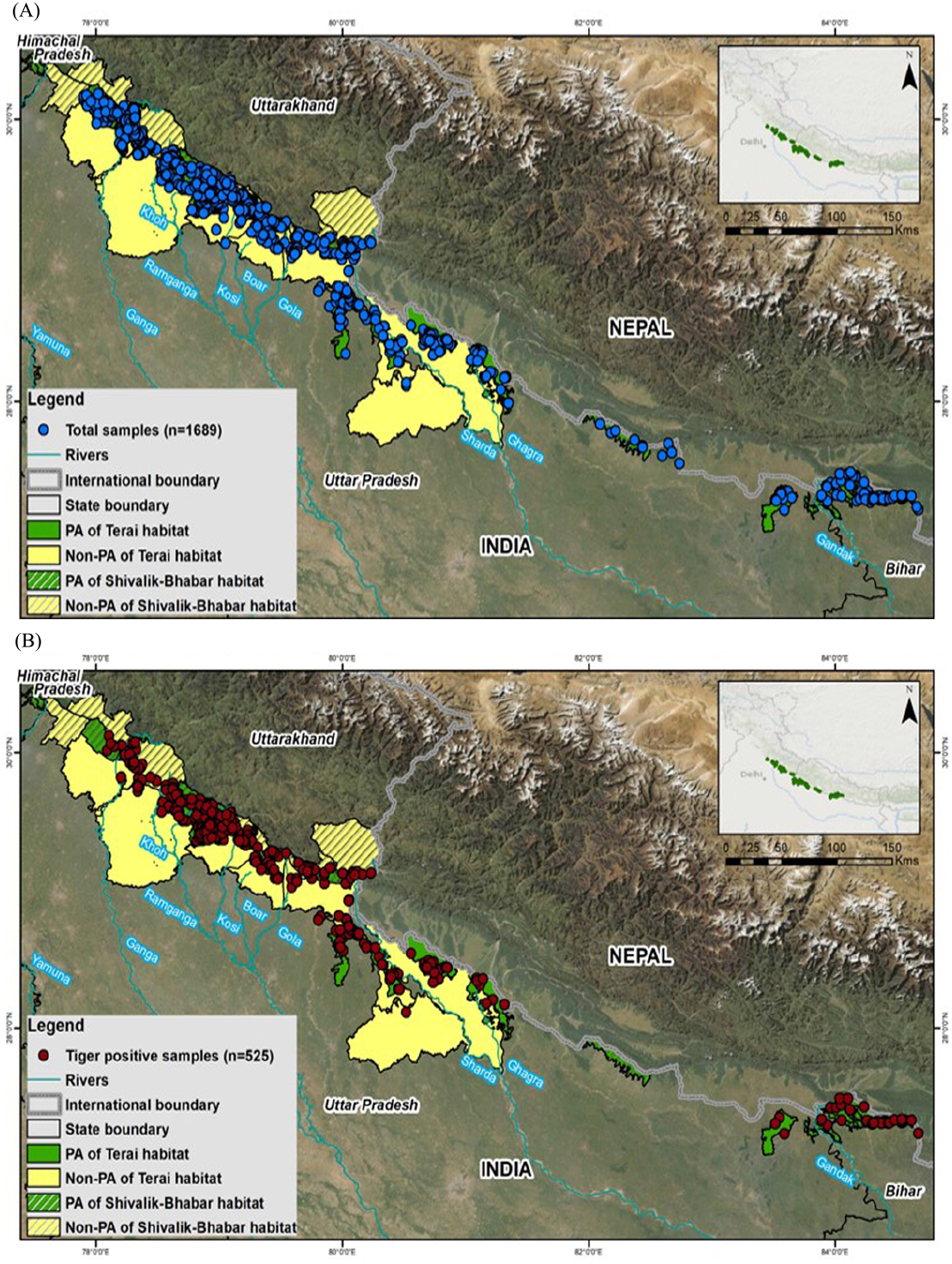

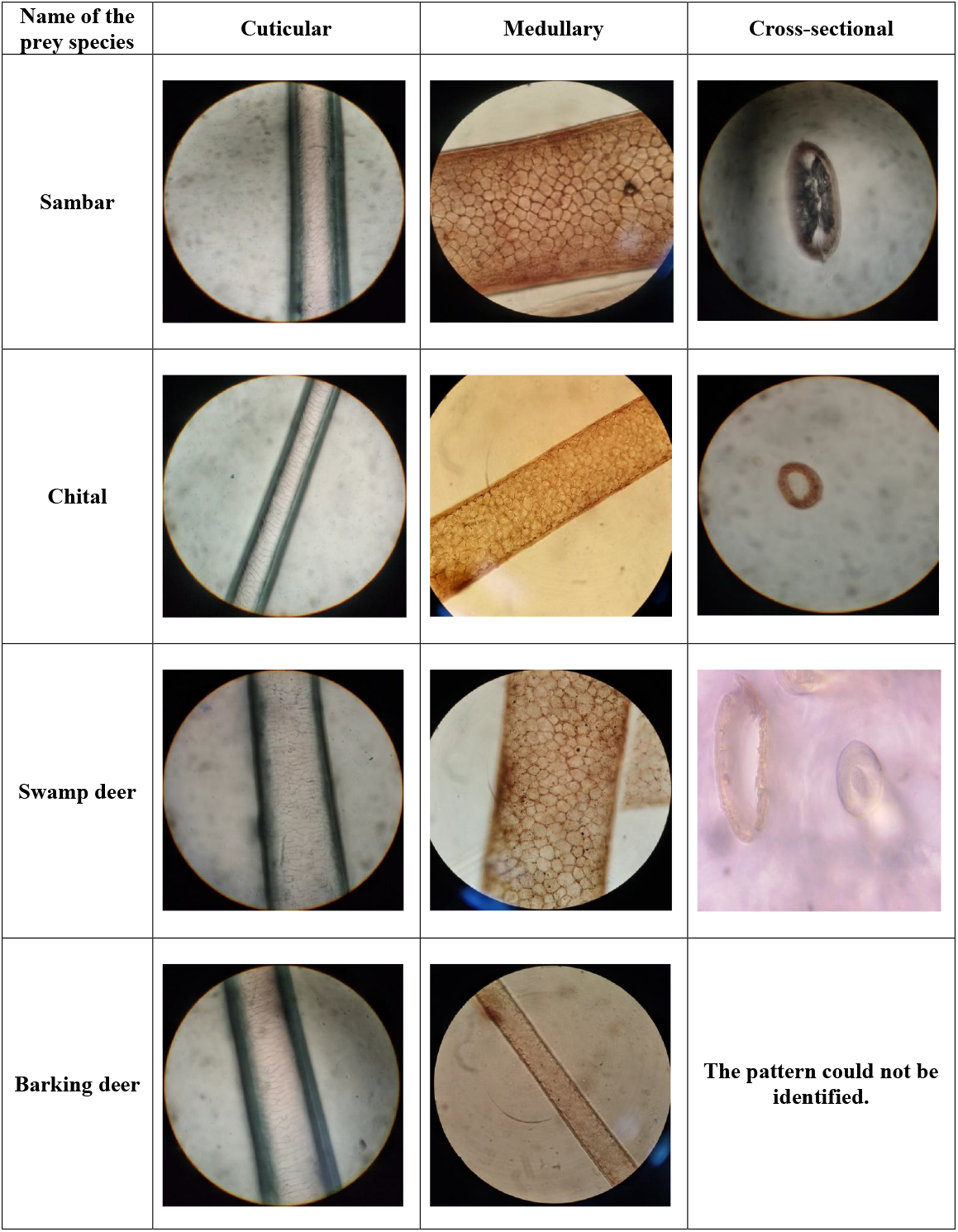

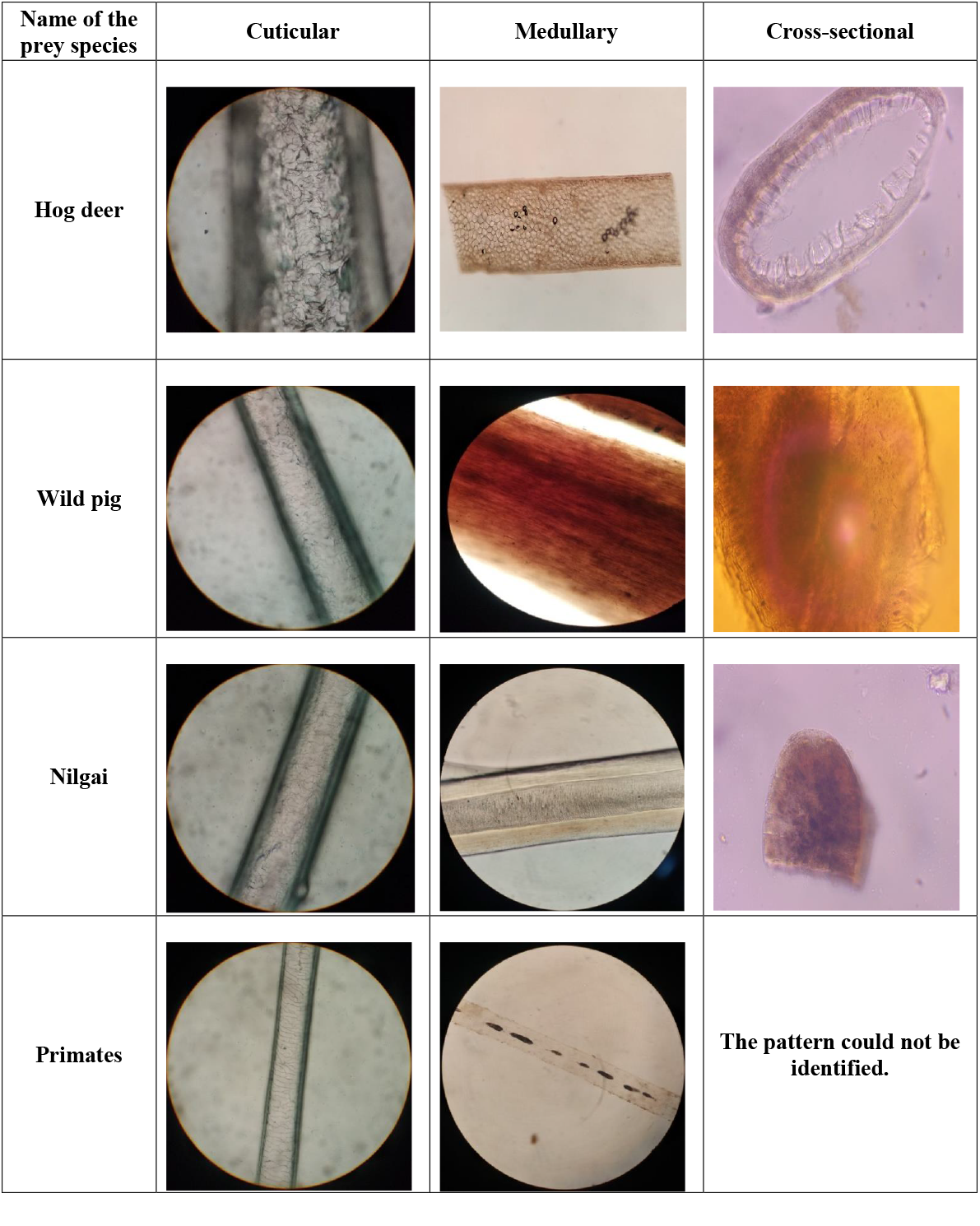

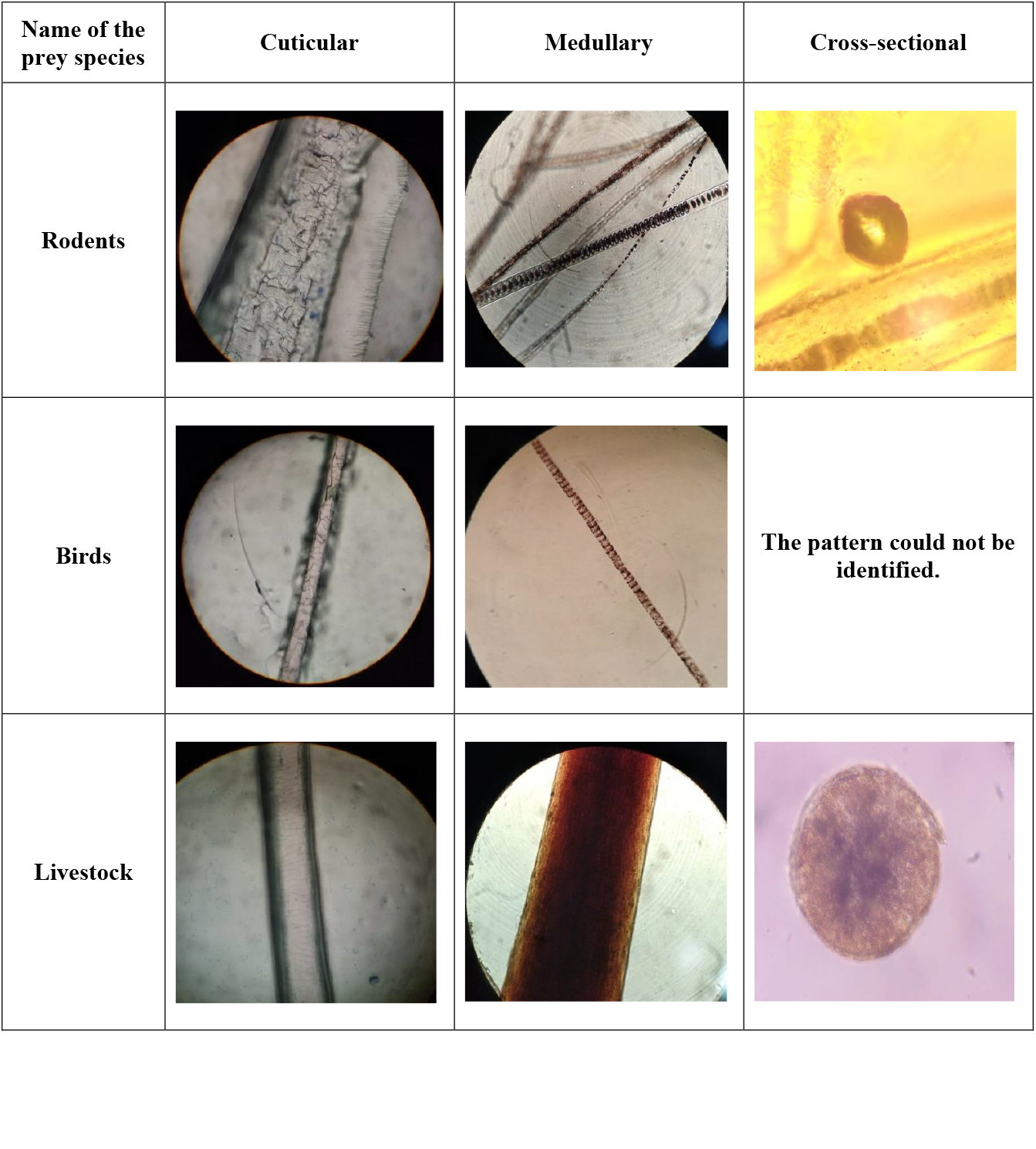

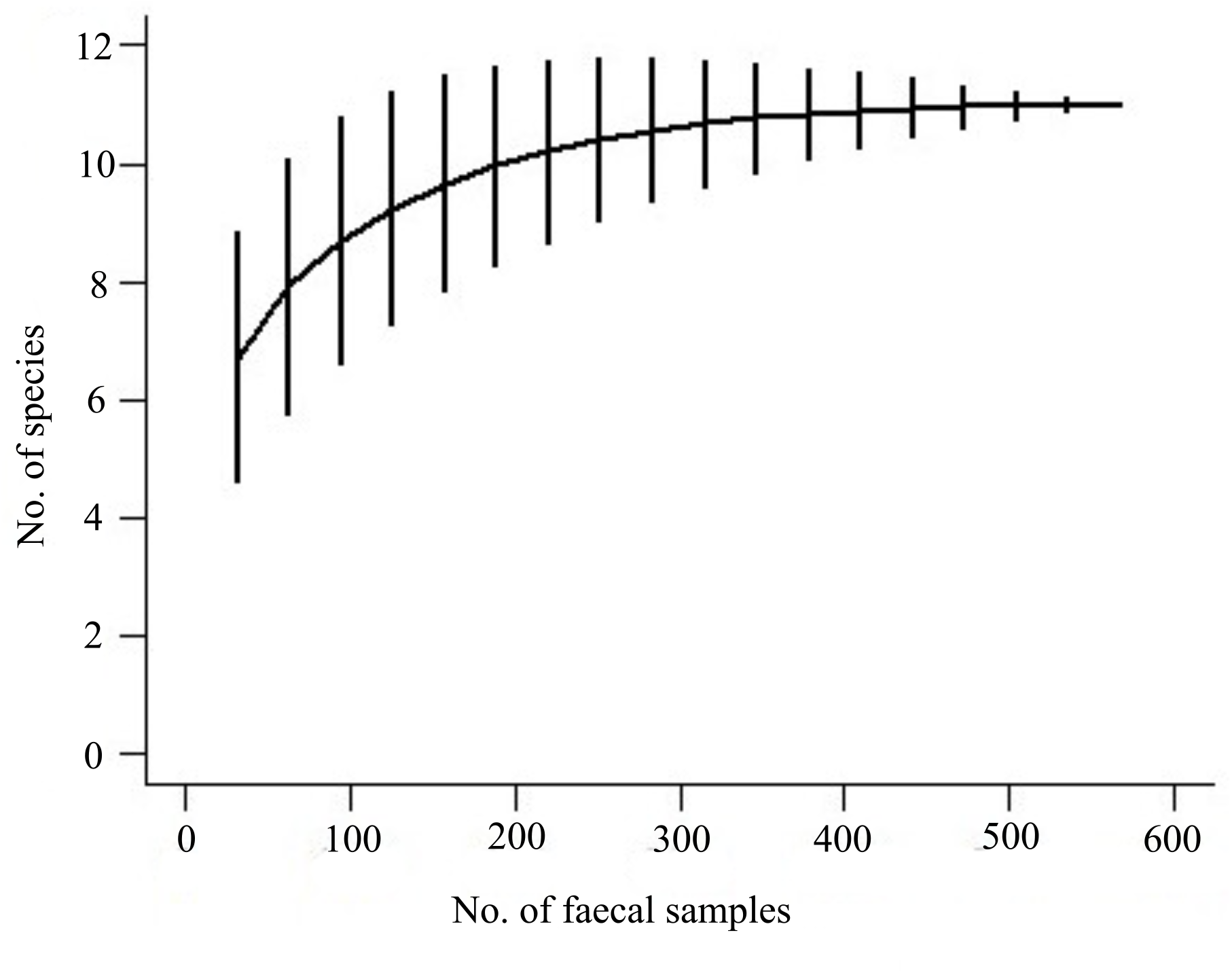

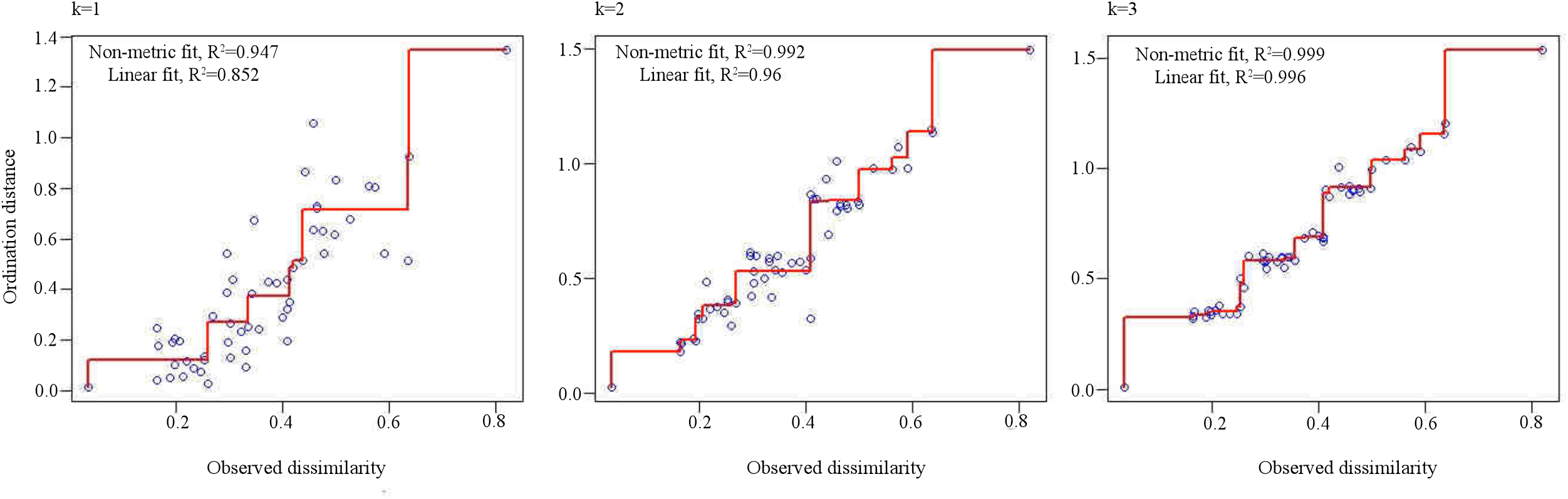

